# The endemic plant species of Bali Ngemba Forest Reserve, Bamenda Highlands Cameroon, with a new Endangered cloud-forest tree species *Vepris onanae* (Rutaceae)

**DOI:** 10.1101/2021.10.06.463416

**Authors:** Martin Cheek, Sebastian Hatt, Jean Michel Onana

## Abstract

We revise and update the records of strict and near-endemic species of the Bali Ngemba Forest Reserve, the largest known surviving patch (c. 8 km^2^ in area) of submontane or cloud forest in the Bamenda Highlands, Cameroon which have lost >96 % of their original forest due to human activities. Nine strict endemics, and 11 near endemics are now documented, a drop from the number recorded after the first survey in 2004, since when five of the provisionally named species have been formally published.

We test the hypothesis that a further one of the provisionally named putative Bali Ngemba new species, *Vepris* sp. A, an 8 – 20 m tall tree from cloud forest in the 1310 – 1600 m altitudinal band, is indeed new to science. We compare it morphologically with other multicarpellate, apocarpous, trifoliolate Cameroon tree species formerly placed in the genus *Oricia* Pierre until they were subsumed into *Vepris* by Mziray (1992). These are *Vepris trifoliolata* (Engl.) Mziray and *V. gabonensis* (Pierre) Mziray. We conclude that *Vepris* sp. A is a new undescribed species here named as *Vepris onanae*. The new species is illustrated, mapped and its conservation status assessed as Endangered using the 2012 IUCN standard due to the threats of habitat clearance from agricultural pressures at its three locations all of which remain formally unprotected.

*Vepris onanae* appears unique among the Guineo-Congolian African oricioid species of *Vepris* in occurring in cloud forest, the other species, apart from *V. renierii* of the Albertine Rift, occurring in lowland forest. It also differs in the very broad, (7.8 –) 11.3 – 18 cm wide leaflets of the flowering stems which have a 6-18(−30) mm long, narrowly triangular acumen (vs leaflets <12 cm wide, acumen absent or short) and in having both subsessile and pedicellate (pedicels 0.25 – 0.3 mm long and 1(– 2) mm long) male flowers (vs male flower pedicels all sessile, or all c. 3 mm long).

We report for the first time on stage-dependent leaf heteromorphy in *Vepris* and characterise a level of sexual dimorphism more advanced than usual in the genus.

We highlight the importance of protecting Bali Ngemba and other forest patches in the Bamenda Highlands if species such as *Vepris onanae* are not soon to become extinct.

## Introduction

As part of a series of studies of *Vepris* by the first and last authors (see below), material from a Cameroonian *Vepris* previously referred to in field studies as *Vepris sp*. A of Bali Ngemba (Harvey *et al*. 2004) was investigated. In this paper we compare the taxon morphologically within *Vepris*, test the hypothesis that this taxon is new to science, formally name it as *Vepris onanae* Cheek.

*Vepris* Comm. ex A. Juss. (Rutaceae-Toddalieae), is a genus with 93 accepted species, 23 in Madagascar and the Comores and 69 in Continental Africa with one species extending to Arabia and another endemic to India (Plants of the World Online, continuously updated). The genus was last revised for tropical Africa by Verdoorn (1926). Founded on the Flore du Cameroun account of Letouzey (1963), eight new species were recently described from Cameroon (Onana & Chevillotte 2015; Cheek *et al*. 2018a; Onana *et al*. 2019; Cheek & Onana 2021), taking the total in Cameroon to 24 species, the highest number for any country globally. The greatest concentration of *Vepris* species in Cameroon is within the Cross-Sanaga Interval (Cheek *et al*. 2001) with 14 species of *Vepris* of which eight are endemic to the Interval. The Cross-Sanaga has the highest species and generic diversity per degree square in tropical Africa including endemic genera such as *Medusandra* Brenan (Peridiscaceae, Breteler *et al*. 2015; Soltis *et al*. 2007). Much of this diversity is associated with the Cameroon Highland areas, different highlands each having a species of a genus e.g. as in *Kupeantha* Cheek (Cheek et al. 2018b). By comparison, neighbouring Gabon has just seven species of *Vepris* (Sosef *et al*. 2006) and just one species, *Vepris lecomteana* (Pierre) Cheek & T. Heller is listed for Congo-Brazzaville (Plants of the World Online, continuously updated), illustrating how under-recorded the Flora of this biodiverse country is, although an endemic species is in the course of publication (Langat *et al*. 2021a). Several Cameroon species are threatened (Onana & Cheek 2011) and in one case considered globally extinct (Cheek *et al*. 2018a), those currently on the IUCN Red List are: *Vepris lecomteana* (Pierre) Cheek & T. Heller (Vulnerable, Cheek 2004), *Vepris trifoliolata* (Eng.) Mziray (Vulnerable, World Conservation Monitoring Centre 1998), *Vepris montisbambutensis* Onana (Critically Endangered, Whittaker & Cheek 2021 and *Vepris adamaouae* Onana (Endangered, Lovell & Cheek 2021). In other parts of Africa species are also highly threatened, e.g., the Critically Endangered *Vepris laurifolia* (Hutch. & Dalziel) O. Lachenaud & Onana of Guinea-Ivory Coast (formerly *V. felicis* Breteler, Cheek 2017; Lachenaud & Onana 2021).

In continental Africa, *Vepris* are easily recognised. They differ from all other Rutaceae because they have digitately (1 –)3(– 5)-foliolate (not pinnate) leaves, and unarmed (not spiny) stems. The genus consists of evergreen shrubs and trees, predominantly of tropical lowland evergreen forest, but with some species extending into submontane forests and some into drier forests and woodland. *Vepris* species are often indicators of good quality, relatively undisturbed evergreen forest since they are not pioneers. New species are steadily coming to light (Cheek *et al*. 2019a).

Species of *Vepris* in Africa extend from South Africa, e.g. *Vepris natalensis* (Sond.) Mziray, to the Guinean woodland in the fringes of the Sahara Desert (*Vepris heterophylla* (Engl.) Letouzey). Mziray (1992) subsumed the genera *Araliopsis* Engl., *Diphasia* Pierre, *Diphasiopsis* Mendonça, *Oricia* Pierre, *Oriciopsis* Engl., *Teclea* Delile, and *Toddaliopsis* Engl. into *Vepris*, although several species were only formally transferred subsequently (e.g. Harris 2000; Gereau 2001; Cheek *et al*. 2009a; Onana & Chevillotte 2015). Mziray’s conclusions were largely confirmed by the molecular phylogenetic studies of Morton (2017) but Morton’s sampling was limited, identifications appeared problematic (several species appear simultaneously in different parts of the phylogenetic trees) and more molecular work would be desirable. Morton studied about 14 taxa of *Vepris*, all from eastern Africa. More recently Appelhans & Wen (2020) focussing on Rutaceae of Madagascar have found that the genus *Ivodea* Capuron is sister to *Vepris* and that a Malagasy *Vepris* is sister to those of Africa. However, the vast majority of the African species including all those of West and Congolian Africa, remain unsampled leaving the possibility open of changes to the topology of the phylogenetic tree when this is addressed.

Characteristics of some of the formerly recognised genera are useful today in grouping species. The “araliopsoid” species have hard, non-fleshy, subglobose, 4-locular fruit with 4 external grooves; the “oriciopsoid” soft, fleshy 4-locular syncarpous fruit; “oricioid” species are 4-locular and apocarpous in fruit; the fruits of “diphasioid” species are laterally compressed in one plane, bilocular and bilobed at the apex; while “tecleoid” species are unilocular in fruit and 1-seeded, lacking external lobes or grooves. There is limited support for these groupings in Morton’s study,

Due to the essential oils distributed in their leaves, and the alkaloids and terpenoids distributed in their roots, bark and leaves, several species of *Vepris* have traditional medicinal value (Burkill 1997). Burkill details the uses, essential oils and alkaloids known from five species in west Africa: *Vepris hiernii* Gereau (as *Diphasia klaineana* Pierre), *Vepris suaveolens* (Engl.) Mziray (as *Teclea suaveolens* Engl.), *Vepris afzelii* (Engl.) Mziray (as *Teclea afzelii* Engl.), *Vepris heterophylla* (Engl.) Letouzey (as *Teclea sudanica* A. Chev.) and *Vepris verdoorniana* (Exell & Mendonça) Mziray (as *Teclea verdoorniana* Exell & Mendonça) (Burkill 1997: 651 – 653). Research into the characterisation and anti-microbial and anti-malarial applications of alkaloid and limonoid compounds in *Vepris* is active and ongoing (e.g., Atangana *et al*. 2017), although sometimes published under generic names no longer in current use, e.g. Wansi *et al*. (2008). Applications include as synergists for insecticides (Langat 2011). Cheplogoi *et al*. (2008) and Imbenzi *et al*. (2014) respectively list 14 and 15 species of *Vepris* that have been studied for such compounds. A review of ethnomedicinal uses, phytochemistry, and pharmacology of the genus *Vepris* was recently published by Ombito *et al*. (2020), listing 213 different secondary compounds, mainly alkaloids and furo- and pyroquinolines, isolated from 32 species of the genus, although the identification of several of the species listed needs checking. However, few of these compounds have been screened for any of their potential applications. Recently, Langat *et al*. (2021b) have published three new acridones and reported multi-layered synergistic anti-microbial activity from *Vepris gossweileri* (I.Verd.)Mziray, recently renamed as *Vepris africana* (Hook.f ex Benth.) Lachenaud & Onana (Lachenaud & Onana 2021).

## Materials and Methods

The taxonomic study is based on herbarium specimens and observations of live material in the Bamenda Highlands and of Mt Kupe in Cameroon. All specimens cited have been seen. The methodology for the surveys in which the specimens were collected is given in Cheek & Cable (1997). Herbarium citations follow Index Herbariorum (Thiers *et al*. continuously updated), nomenclature follows Turland *et al*. (2018) and binomial authorities follow IPNI (continuously updated). Material of the new species was compared morphologically with material of all other African *Vepris*, principally at K, but also using material and images from BM, EA, BR, FHO, G, GC, HNG, P and YA. Specimens at WAG were viewed on the Naturalis website (https://bioportal.naturalis.nl/). The main online herbarium used during the study apart from that of WAG was that of P (https://science.mnhn.fr/all/search). Herbarium material was examined with a Leica Wild M8 dissecting binocular microscope fitted with an eyepiece graticule measuring in units of 0.025 mm at maximum magnification. The drawing was made with the same equipment using a Leica 308700 camera lucida attachment.

For the extinction risk assessment, points were georeferenced using locality information from herbarium specimens. The map was made using simplemappr (Shorthouse 2010). The conservation assessment was made using the categories and criteria of IUCN (2012), EOO was calculated with GeoCat (Bachman *et al*. 2011).

## Results

The earliest known specimen collected of *Vepris* sp. A. was *Etuge* 1814 made in March 1996 from Mt Kupe. It was initially identified as *V. gabonensis*, then as being more likely *V. trifoliolata* (with the statement that more material was needed). This specimen was cited with qualifications as the last species in the Checklist for Mwanenguba, Bakossi and Mt Kupe (Cheek *et al*. 2004: 396). Indeed *Vepris* ap. A shares many features with both these species which were formerly placed together in the genus *Oricia* together with *Vepris suaveolens* and *V. lecomteana* (Letouzey 1963). These five central African taxa all occur in Cameroon and share a female flower with four separate carpels which however are closely appressed to each other, appearing as a single subglobose structure, the separation concealed by dense tomentum, and they are united by their joined, common, 4-lobed, flat, black, glabrous stigma. In fruit at least one carpel aborts so that only 1 – 3 mericarps mature, the remainder appearing vestigial. This group of species have stems, often other vegetative structures, and inflorescences usually densely hairy, unusual in a genus which is usually glabrous, or nearly so. Additional oricioid species occur in the montane forest of Rwanda (*V. renierii* C.C.G.Gilbert) and *V. bachmannii* (Engl.)Mziray (glabrous, unusually) from Mozambique to Kwa-Zulu-Natal in S Africa.

*Vepris* sp. A is unlikely to be confused with *V. lecomteana* since that species bears 5, not 3 leaflets, and these leaflets are sessile not petiolulate. Nor is confusion likely with *Vepris suaveolens* since that species has leaflets only 1.5 – 10 cm broad (vs 8.7 – 14.8 cm) with an obtuse or acute apex (vs distinctly acuminate), and the flowers are all sessile (vs at least some flowers pedicellate). Potential confusion with *Vepris gabonensis* and *V. trifoliolata* can be resolved by referring to table 1, below.

**Table 1.**
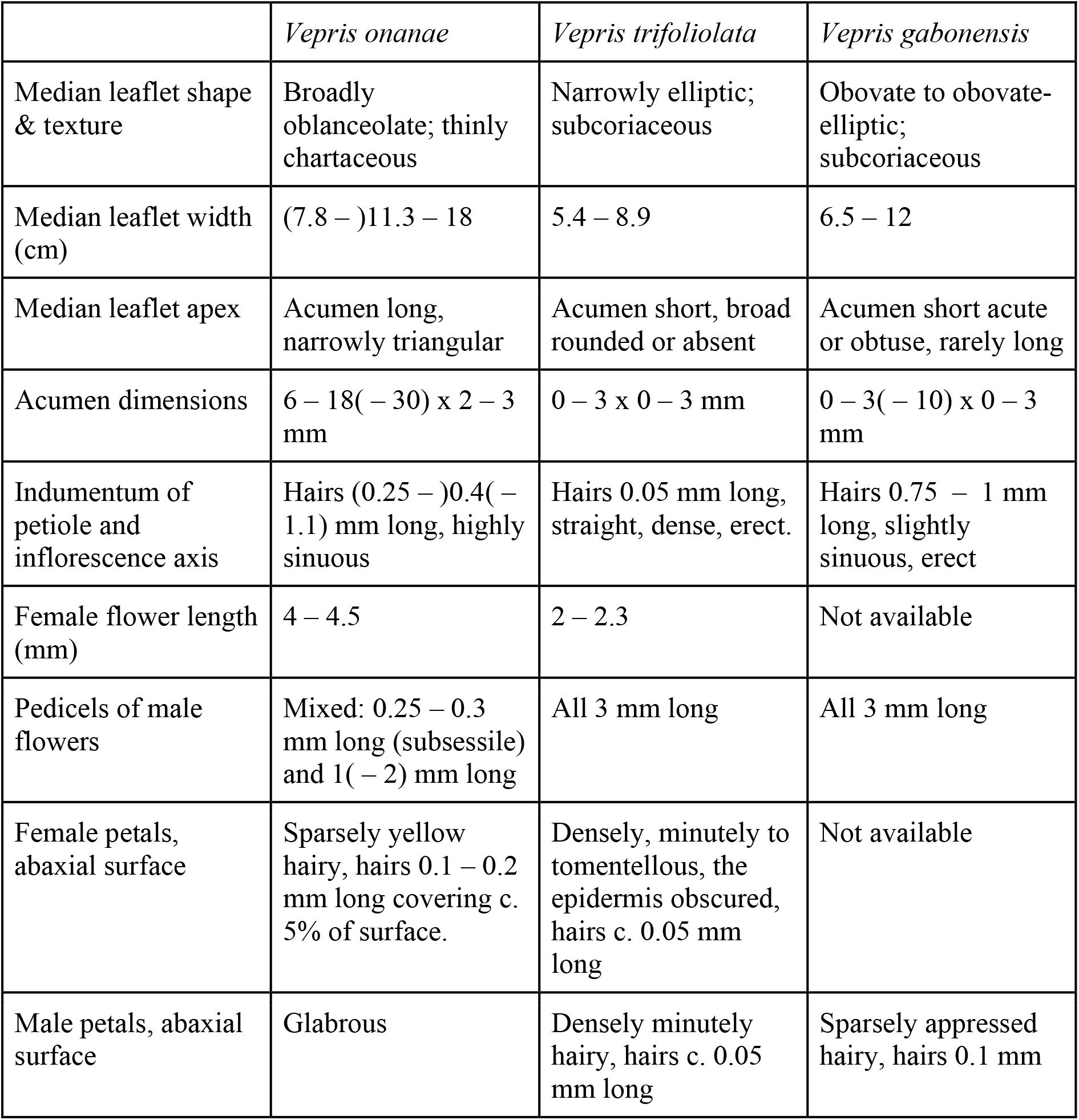
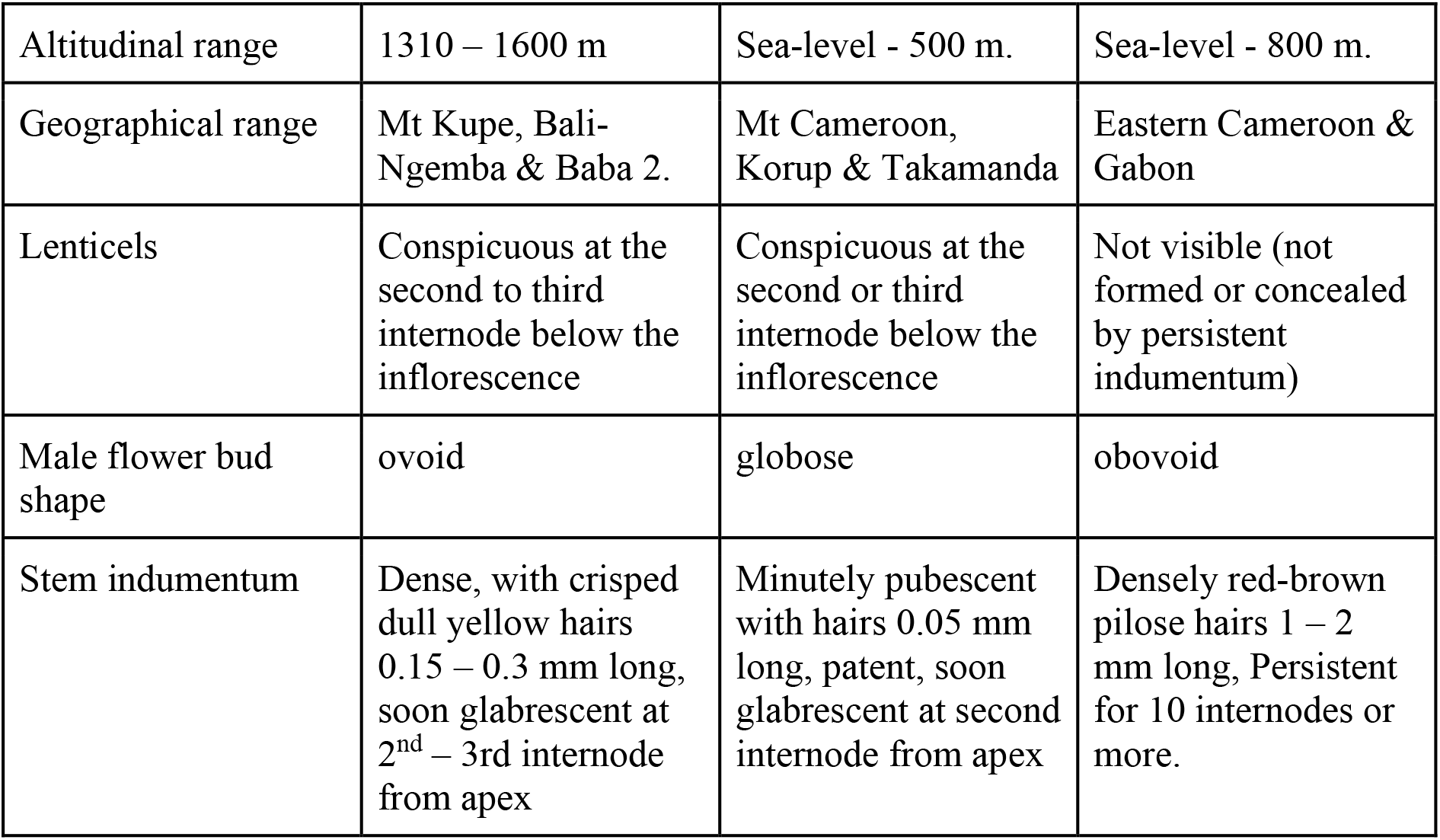
Diagnostic characters separating *Vepris onanae* from *V. trifoliata* and *V. gabonensis*

### Vepris onanae

Cheek sp. nov. Fig. 1. Type: Cameroon, North West Region, Mezam Division, Baba 2 village community forest, 1610 m alt., female fl. 12 April 2002, *Onana* 2006 with F. Nana, Ela Nangmo, P.Nkankano, Bakary, Malota & Njume (holotype K000593330; isotype YA).

**Fig. 1.**
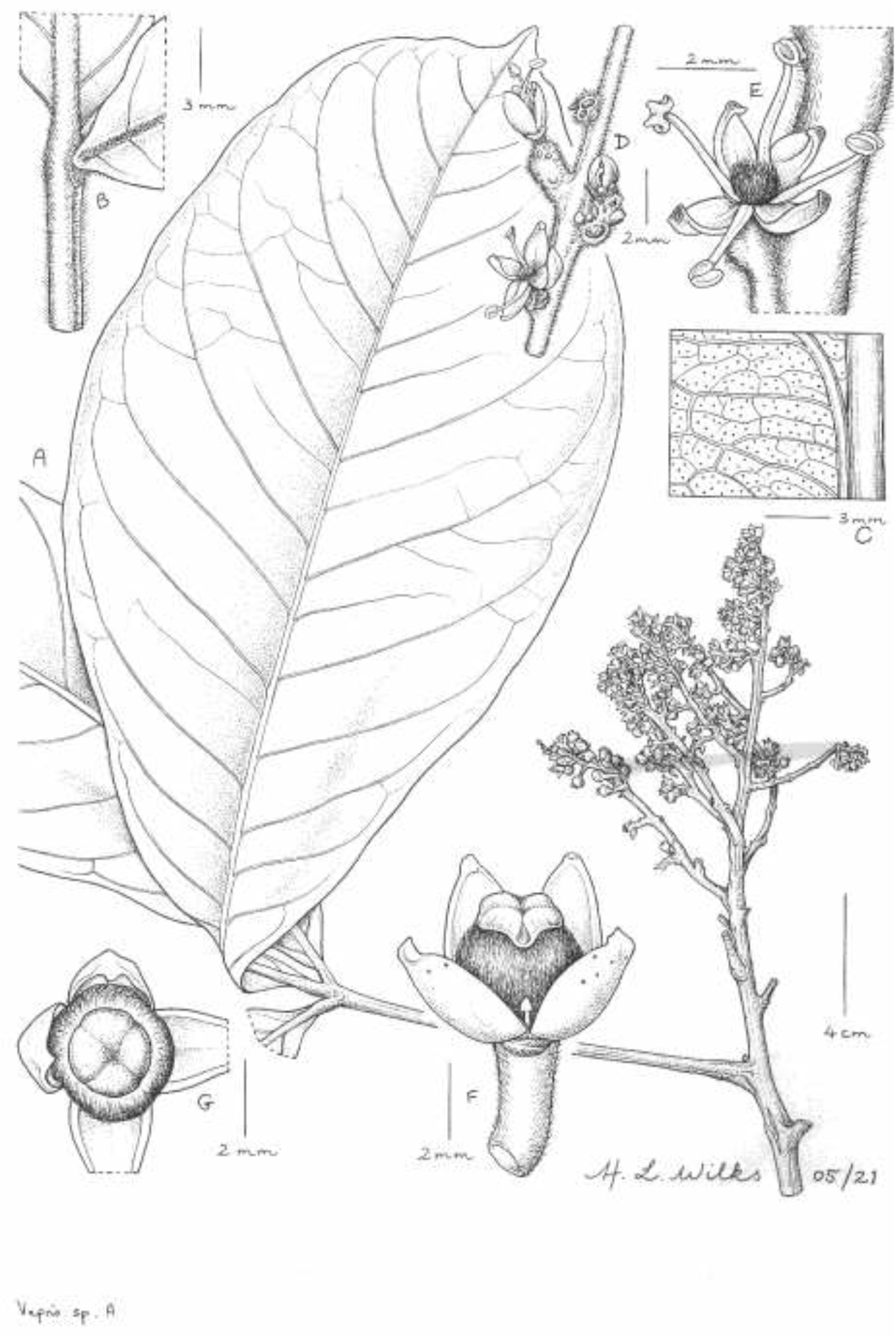
*Vepris onanae*. **A** habit, flowering stem with female inflorescence; **B** base of leaflets and indumentum; **C** gland dots on abaxial leaf surface; **D** portion of male partial inflorescence; **E** male flower, showing sessile attachment; **F** female flower, side view, showing vestigial stamen and oil glands; **G** female flower, top view, with two petals pulled back. **A-C, F** & **G** from *Onana* 2006, **D** & **E** from *Zapfack* 2039. DRAWN BY HAZEL WILKS.

Syn. *Vepris* sp. A Cheek (Harvey *et al*. 2004: 124)

*Vepris trifoliolata sensu* Cheek *et al*. (2004:396 quoad *Etuge* 1814) non (Engl.)Mziray

*Dioecious* evergreen tree 8 – 20 m tall. Trunk brown, cylindric, sapwood white. Leafy stems terete, (3.5 –)5 – 7 mm diam., internodes 1.7 – 2.9(– 6.7) cm long, densely woolly-tomentose, hairs simple, 0.1 – 1 mm long, of two classes, longer hairs (c. 1 mm long) often crispate, patent, pale golden-brown, at stem apex covering c.100% of the surface, gradually glabrescent, decreasing in density with distance from apex, c. 50% cover at third node from apex, exposing the shorter (c. 0.1 mm long), straight, closely appressed hairs; older internodes glabrous, epidermis drying pale grey to yellow-green, longitudinally fluted, becoming densely lenticellate, lenticels white, slightly raised, shortly elliptic or orbicular, 0.5-0.75 mm long *Leaf* phyllotaxy alternate, trifoliolate, drying mid green above, pale green below, 18 – 50 cm long, median leaflet elliptic-oblanceolate, (14.5 –)21 – 36.1 × (4 –)8.7 – 14.8(– 17.4) cm, lateral leaflets 70 – 85% the length of the median, elliptic-lanceolate, apex distinctly acuminate/narrowly-acuminate, acumen 0.7 – 1.8(– 3.1) × 0.5 – 0.9(– 1.8) cm, base of leaflet cuneate; margin slightly thickened, revolute; midrib and secondary nerves, raised, thickened, secondary nerves, 9 – 12(– 14) on each side of the midrib, yellow-green, arising at c. 60° from the midrib, arching towards the leaflet apex, forming a looping inframarginal nerve c. 2 mm from the margin, glabrescent. Intersecondary nerves 1(– 2), arising c. 80° from the midrib, extending towards margin before branching. Tertiary and quaternary nerves inconspicuous but visible, very faintly raised; essential oil glands 4 – 8(– 13) per mm^2^, orbicular, 0.1 – 0.15 mm diam., appearing black, often indistinct on the abaxial surface (Fig. 1C), invisible on adaxial surface and colourless in transmitted light; midrib tomentose abaxially, adaxially broad, slightly sunken, densely shortly hairy (Fig. 1B), otherwise glabrous, petiolules pulvinate proximally (larger leaves), distally adaxially flat, margins inconspicuously narrowly winged, decurrent from blade, revolute, (0.1 –)0.4 – 0.8(– 1.3) cm long, tomentose with dense simple, crispate, golden hairs c. 0.2 mm long. Petiole articulated with the petiolules, terete, longitudinally fluted, sometimes with 2 indistinct low ridges, (3 –)9.2 – 14 cm long, 2.5 – 4 mm diam., hairs scattered, golden, c 0.15 mm long, spreading to appressed, glabrescent.

*Male inflorescences* terminal, or axillary in distal axils, paniculate, 14.9 – 15.5 x 8 – 27 cm, c. 100-flowered, peduncle 0.7 – 3.5 cm long; rachis internodes 15 – 17, 0.6 – 2 cm long; partial inflorescences 4 – 15 cm long, bracts caducous, partial-peduncles alternate, 0.7 – 2 cm long, second order branches 3-40 mm long, becoming shorter distally, internodes 1 – 8 mm long, bearing third order branches reduced to 1-2 mm long, each bearing single flowers or 2 – 4-flowered glomerules (Fig. 1D), glomerulous flowers subsessile, pedicels 0.2-0.3 mm long, single flower pedicels 1(−2) mm long; bracteoles subtending glomerules narrowly triangular, 0.5 mm long, densely hairy. *Female inflorescences* (Fig. 1A) terminal 11 × 10.5 cm, c. 100- flowered, peduncle 1.2 cm long; rachis internodes c. 8, 0.5 – 1.4 cm long; partial-inflorescences 4.5 – 8 cm long, partial-peduncles alternate, 1 – 1.6 cm long, bracts caducous, bracteoles caducous or absent. Second order branches absent, flowers single, or in pairs. Pedicels 3 – 6 mm long, scattered with dense golden indumentum. *Male Flowers* (Fig. 1E) greenish white, c. 3 mm wide. Sepals 4, broadly ovate, slightly spreading and concave, c. 0.5 × 0.5 mm, apex acute to obtuse, short golden hairs near base of sepal, margins sometimes sparsely fimbriate. Petals 4, ovate to elliptic-oblong, 2 – 2.2 × 1 – 1.3 mm, apex acute-obtuse, often incurved near apex to form small hook-like appendage, glabrous; oil glands not visible. Stamens 4, equal, always exceeding petals, filaments 2.2 – 3 mm long, ribbon-like; anthers yellow, ellipsoid, 0.6 – 0.75 × 0.4 mm. Pistillode (vestigial ovary), broadly ovoid, c. 0.75 × 0.75 mm, obscured by dense crispate golden hairs, c 0.4 mm long covering c. 95% of the surface. *Female Flowers* (Fig. 1F&G) white, c. 4 mm long, 5 – 9 mm wide. Sepals 4, broadly ovate, slightly spreading and concave, c. 1 × 1.1 mm, apex acute, covered in moderately dense short golden hairs, with highest density at the base. Petals 4, broadly ovate, 3.5 – 4 × (1.5 –)2.5 mm, apex acute, often incurved near apex to form small hood, lacking appendages, abaxial surface moderately to sparsely covered in short, thick, appressed, golden hairs 0.1 – 0.2 mm long; oil glands orange to dark brown, the larger glands dark brown, elliptic or orbicular, 0.6 – 0.8 mm diam., 8 – 12 scattered on abaxial and adaxial surfaces but concentrated in the distal half. Ovary apocarpous, carpels 4, coherent, appearing 4-lobed, subglobose, 1.75 – 2.5 × 2-3 mm, densely yellow tomentose, hairs spreading, c. 1 mm long; stigma 4-lobed or square, 1.5-2 mm diam., flat, margins revolute, dark brown to black, glabrous,; stamens (vestigial), 0 – 4, ribbon-like, 1 – 3 mm long, anther deltate-sagittate, 0.4 mm long. Fruit not seen.

### RECOGNITION

Similar to *Vepris trifoliolata* (Engl.)Mziray, differing in larger (7.8 –)11.3 – 18 cm wide, broadly oblanceolate median leaflets (vs 5.4 – 8.9 cm wide, narrowly elliptic), with a long, triangular acumen (vs acumen barely developed, short and broad), indumentum of long sinuous, sometimes branched hairs (0.1 – 1.1 mm long (vs short straight hairs c. 0.05 mm long), male flower buds ovoid-ellipsoid, petals glabrous abaxially, pedicels 0.25 – 0.3 mm and 1(– 2) mm long (vs globose, puberulent, 3 mm long).

### DISTRIBUTION

Cameroon, South West Region, Mt Kupe and North West Region, Bali-Ngemba & Baba 2 forests. Map 1.

**Map 1.**
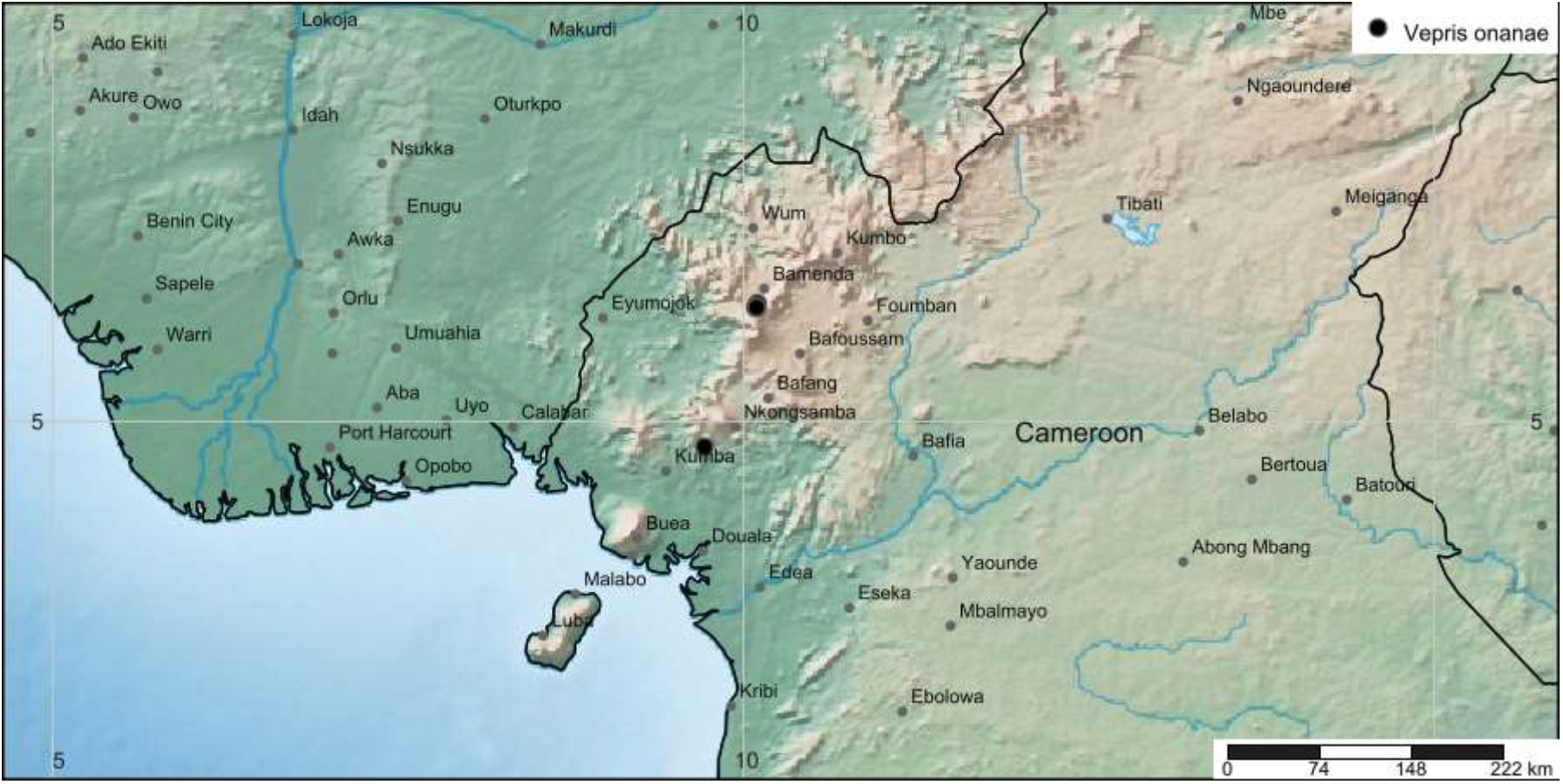
Global distribution of *Vepris onanae*

### SPECIMENS EXAMINED. CAMEROON

**North West Region, Mezam Division, Bali, Bali Ngemba Forest Reserve**, Mantum, First Valley, 1400 m alt., st. 11 July 2000, *Cheek* 10437 with Wultoff & R. Ngolan (K000593334, YA); ibid. Kup Fujah Forest along main trail N from Mantum, male fl. 12 Apr. 2004, *Darbyshire* 385 with Nukam, Barlas, de Martrin, Nana and Atem (K, MO n.v., SCA n.v., WAG n.v., YA n.v.); ibid. Near stream, 1400 m alt., st. 9 Nov. 2000, *Ghogue* 1048 with Ayako, Kate, Julia, Doh Lawan (K00593332, MO n.v., WAG n.v., YA n.v.); 1.5 miles along Cypress way, 1450 m alt., male fl. 10 April 2002, *Pollard* 930 (K00593328, YA n.v.) River Forest, 1310 m alt., male fl. 11 April, 2002, *Zapfack* 2039 with Asase, Kom, Orurke, Azamah, Oketch, Nana V. (K000593333, SCA n.v., YA n.v.); **Baba 2 village community forest**, 1610 m alt., st. 6 Oct 2001, *Cheek* with Etuge, Njume, Rowe, Rengeana, Nkeng & Ngolan 10747 (K000593331, YA); ibid, female fl. 12 April 2002, *Onana* 2006 (K000593330, YA). **South West Region, Kupe-Muanenguba Division**, Nyassoso, Mt Kupe, 1600 m alt., male fl. 26 March 1996, *Etuge* 1814 (K000198269, YA n.v.).

### HABITAT

Lower submontane forest with *Pterygota mildbraedii* Engl. as emergent trees from canopy (North West Region only). Associated species of tree are *Allanblackia gabonensis* (Pellegr.)Bamps (Clusiaceae), *Alsophila* spp. (Cyatheaceae), *Noronhia africana* (*Knobl*.) *Hong-Wa & Besnard* (*Chionanthus africanus* (Knobl.) Stearn) (Oleaceae), *Macaranga occidentalis* (Müll.Arg.) Müll.Arg. (Euphorbiaceae), *Polyscias fulva* (Hiern) Harms (Araliaceae), *Pseudagrostistachys africana* (Müll.Arg.) Pax & K.Hoffm. (Euphorbiaceae), *Santiria trimera* (Oliv.) Aubrév. (Burseraceae), *Trichilia dregeana* Sond. (Meliaceae) (Harvey *et al*. 2004:17 – 18) alt. 1310 – 1600 m.

### CONSERVATION STATUS

*Vepris onanae* is known only from the neighbouring and formerly contiguous forests of Bali Ngemba and Baba 2, and from a solitary specimen from the 100 km distant Mt Kupe. Bali Ngemba is a government managed Forest Reserve intended for timber production. However, for several years it has been reported that staffing of forest guards has been insufficient to protect it, and incursions have occurred, removing trees for timber and clearing forest understorey for cultivation of crops (Kenneth Tah pers. comm. to Cheek Feb. 2021). The forest at Baba 2 belongs and is managed by the community of that name and produces ‘Cola nuts’ from the trees of the indigenous *Cola anomala* K. Schum. The canopy is more intact than that of Bali Ngemba, but there is a risk that enrichment planting of *Cola* may be negative for the population of *Vepris onanae*. At Mt Kupe it is extremely rare, with only a single collection at its sole location on the main path to the summit from Nyasoso despite intensive general collection effort of all plants over several years. Forest clearance upslope for agriculture has reached the altitude at which it occurs. It cannot be ruled out that *Vepris onanae* will be found at additional locations outside of its present range. Yet, many thousands of specimens have been collected at points of the compass around Bali Ngemba, Baba 2 and Mt Kupe in other surveys, despite which this species has not been found (Cheek 1992; Cheek *et al*. 1996; Cable & Cheek 1998; Cheek *et al*. 2000; Chapman & Chapman 2001; Cheek *et al*. 2006; Cheek *et al*. 2010; Harvey *et al*. 2010; Cheek *et al*. 2011). The extent of occurrence is calculated as 44 km^2^ and the area of occupation as 12 km^2^ using 4km^2^ grid cells preferred by IUCN. Therefore, we here assess *Vepris onanae* as EN B1ab(iii) + B2ab(iii). The species might be considered severely fragmented since the three forests in which it survives have become separated from each other due to clearance for agriculture. Evidence for seed dispersal by primates (chimpanzees) was recently presented for another new species of *Vepris* (Langat *et al*. 2021a) for which traversal of human created habitat is known to be problematic (Cheek et al. 2021a). While the second *Vepris* species at Bali-Ngemba, *Vepris bali* occurred at higher altitude and is considered likely to be extinct (Cheek *et al*. 2018a), we consider it likely that *Vepris onanae* survives there at present since it was found to be relatively frequent in our surveys of 2000 – 2004, witness the number of specimens cited. However, unless incursions at Bali Ngemba are stopped, it is only a matter of time before it also becomes extinct there.

### PHENOLOGY

Flowering March and April; fruiting unknown; sterile in July, Oct., Nov.

### ETYMOLOGY

Named in honour of Dr Jean-Michel Onana, Lecturer in Botany at University of Yaoundé I, Cameroon, champion of plant conservation in Cameroon, specialist in Sapindales (Burseraceae, author of Flore du Cameroun Burseraceae (Onana 2017), co-chair of the IUCN Central African Red List Authority for Plants, former Head of the Biodiversity Program of IRAD and the National Herbarium of Cameroon (2005–2016), author of the Taxonomic Checklist of the Vascular Plants of Cameroon (Onana 2011), a synopsis of the endemic and rare vascular plants of Cameroon (Onana 2013) and co-author of the Red Data Book of the Plants of Cameroon (Onana & Cheek 2011). He led field teams of YA staff working with those of K that resulted in the collection of numerous specimens during these surveys and personally collected the type specimen and only known female specimen of this species in the field (*Onana* 2006, K, YA). *Ledermanniella onanae* Cheek (Cheek 2003), *Diospyros onanae* Gosline (Gosline 2009), *Psychotria onanae* O.Lachenaud (Lachenaud 2019) and *Deinbollia onanae* Cheek (Cheek *et al*. 2021a) are also named for him.

### VERNACULAR NAMES & USES

None are recorded.

### NOTES

Bali Ngemba and Baba 2 in the Bamenda Highlands, are the centre of distribution of this tree, with seven of the eight collections. That the species should be absent from Mt Oku and the Ijim Ridge Forest which is well known for its threatened plant species (Cheek *et al*. 2000, Maisels *et al*. 2000) is because this location contains only montane forest occurring at 2000 m alt. or higher, above the range of this species. This also accounts for its absence at Dom (Cheek *et al*. 2010) which is also higher altitude than the range of *Vepris onanae*, but not at Lebialem Highlands (Harvey *et al*. 2010) (now the Tofala Hills Sanctuary), where it may be present but remain uncollected since this location is intermediate between Bali Ngemba and Mt Kupe, over 100 km to the south whence it was collected in the same altitudinal band as in the Bamenda Highlands.

## Discussion

### Sexual dimorphism in *Vepris onanae*

Previous studies of *Vepris* (see introduction) have not covered the subject of sexual dimorphism in the genus apart from the obvious fact that male flowers have fully formed, functional stamens, but reduced, vestigial ovaries, pistillodes. In contrast female flowers have fully formed ovaries but reduced stamens, bearing minute non-pollen bearing anthers, staminodes. However, in *Vepris onanae*, additional morphological disparities occur between the four male specimens and the one female, especially in the structure of the inflorescence. While in the female the principal axis of the inflorescence bears partial-inflorescences bearing single or paired flowers along their length (Fig. 1A), in the male, the partial-inflorescences produce two further orders of branches, and most of the flowers are borne in 2 – 5-flowered glomerules (Fig. 1D), additionally the female flowers are borne on pedicels 3 – 6 mm long, and the petals abaxially hairy, while the glomerulous male flowers have the abaxial surfaces of the petals glabrous and the flowers are subsessile, and those borne singly have 1(– 2) mm long pedicels. The sample size available is so low that we cannot conclude that the 4:1 male: female ratio seen is representative or not, but it does suggest that male individuals might outnumber females. A similar scenario is seen in the Burseraceous genera *Canarium* and *Dacryodes* (Onana 2017).

### Stage-dependent leaf heteromorphy in *Vepris*

Most of the specimens seen of *Vepris onanae* were collected in flower in April-July, in the early wet season. However, *Ghogue* 1048(K) was collected in early November, and was sterile. It differs from the other specimens in having at the stem apex 3 – 5 highly reduced leaves each 1 – 2 cm long, forming a loose bud. Since November is the beginning of the dry season, this may represent the resting bud of a dormant stage. The 2 – 3 leaves subtending this structure are short, only 7 – 8 cm long compared with those produced earlier in the season when the median leaflets alone are 21 – 36 cm long. The same pattern with similar chronology is also seen in species of the genus *Cola* Schott & Endl. which grow in the same habitat (Cheek et al. (2020a). Such stage dependent leaf heteromorphy has not been reported before in *Vepris* to our knowledge.

### The Cameroon Highlands, Bamenda Highlands and Bali Ngemba

The Bamenda Highlands are the section of the Cameroon Highland chain that extend through North West Region, Cameroon. Much of the chain at this point is relatively flat and fertile, at 2000 m altitude, and is referred to as the “High Lava Plateau” by Hawkins & Brunt (1965). It has a lower rainfall, often less than 2 m per annum, in comparison to the southern section that adjoins it and continues southwards to the sea at Mt Cameroon, rainfall here reaching over 3 m per annum, and over 10 m per annum at the foot of Mt Cameroon (Cable & Cheek 1998; Courade 1974). The Bamenda Highlands are densely inhabited and the original forest vegetation has been almost entirely replaced through clearance by secondary grassland, sometimes invaded by secondary woodland (Hawkins & Brunt 1965). For this reason, the area is known as “The Grassfields”.

Those few areas that remain under indigenous trees are mainly sacred forests, reserved for traditional ceremonies by the tribal communities that manage them. The largest surviving forest area, at c.200 km^2^, is Mt Oku (Kilum-Ijim), at 3011 m the highest point in the Bamenda Highlands however this forest is all above 2000 m alt., i.e. montane, forest below that altitude being almost entirely lost. It has been calculated that c. 96.5 % of the original forest in the Bamenda Highlands has been lost (Cheek *et al*. 2000).

### The endemic plants of Bali Ngemba

Within the Bamenda Highlands, Mt Oku (Kilum-Ijim) has been well-known for the importance of its biodiversity for both plants and animals for decades (Thomas 1986; Collar & Stuart 1988; Tame & Asonganyi 1995; Cheek *et al*. 1997; Maisels *et al*. 2000; Cheek *et al*. 2000). In contrast, Bali Ngemba Forest Reserve was almost unknown until we began our surveys there in 2000 – 2004 resulting in the Plants of Bali Ngemba Forest reserve (Harvey *et al*. 2004). After the collection of 1381 herbarium specimens in this period, and subsequent identifications, among the hundreds of plant species recorded for the first time, 38 threatened species were recorded, apart from a further 24 that were considered new to science and which were expected also to be assessed as threatened once formally published. In total 12 species were considered to be unique (endemic) to Bali Ngemba, with a further 11 near-endemic (Harvey *et al*. 2004). Most of the species do not occur at Mt Oku because of the differing altitudinal ranges, Bali Ngemba being submontane forest (c. 1300 – 2000 m alt.)

Many of the endemics were known under provisional names only. Since that time several of these have been formally published. Some of the species previously thought to be endemic have been discovered at additional locations so that they are no longer either endemic to Mt Bali Ngemba (known only from Bali Ngemba) or near-endemic (here defined as from Bali Ngemba and one or two other locations).

Examples of species previously considered endemic but no longer so are *Monanthotaxis* sp. of Bali Ngemba soon to be published as *M. vulcanica* P.H.Hoekstra (Couvreur *et al*. 2021) now recorded also from three other locations near to the Bamenda Highlands. Similarly, *Tricalysia* sp. aff. *ferorum* is now published as *T. elmar* Cheek and is known also from Mt Kupe, Ngovayang and the Rumpi Hills (Cheek *et al*. 2020b; Lachenaud pers. comm. to Cheek 2019). One taxon thought to be new to science has subsequently been reidentified, namely *Psychotria* sp. A aff. calva as *P. subpunctata* Hiern (Lachenaud 2019). Others new to science have proved to be widespread through the Cameroon Highlands, such as *Deinbollia* sp. 1 (now *Deinbollia oreophila* Cheek (Cheek & Etuge 2009a).

Below, in Table 2, we update the known endemics of Bali Ngemba. In addition to those species originally documented as such in Harvey *et al*. (2004) often now under other names, we include a species that had not been included there as an endemic, the near endemic *Scleria afroreflexa* Lye (Lye & Pollard 2003).

**Table 2.**
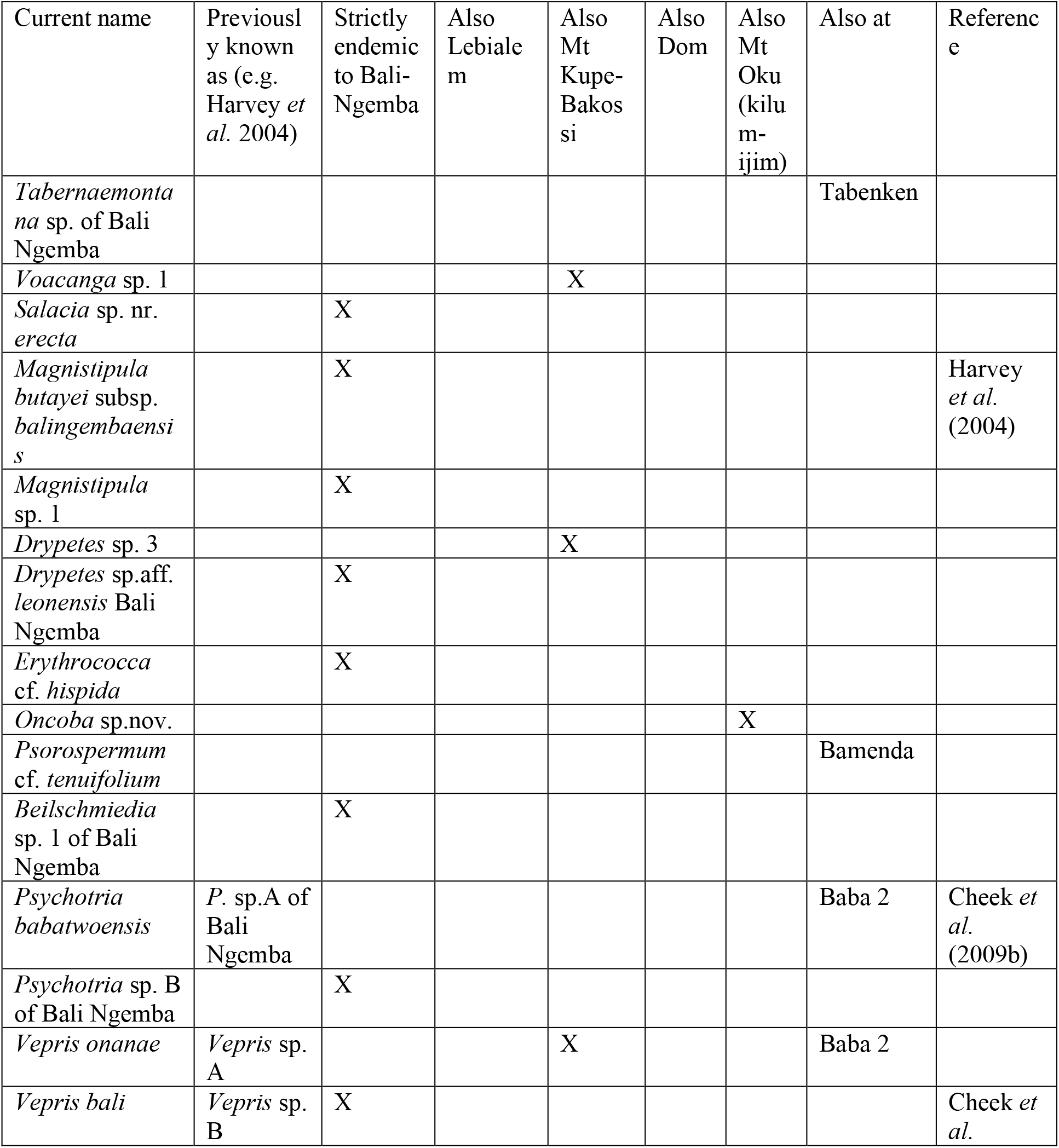
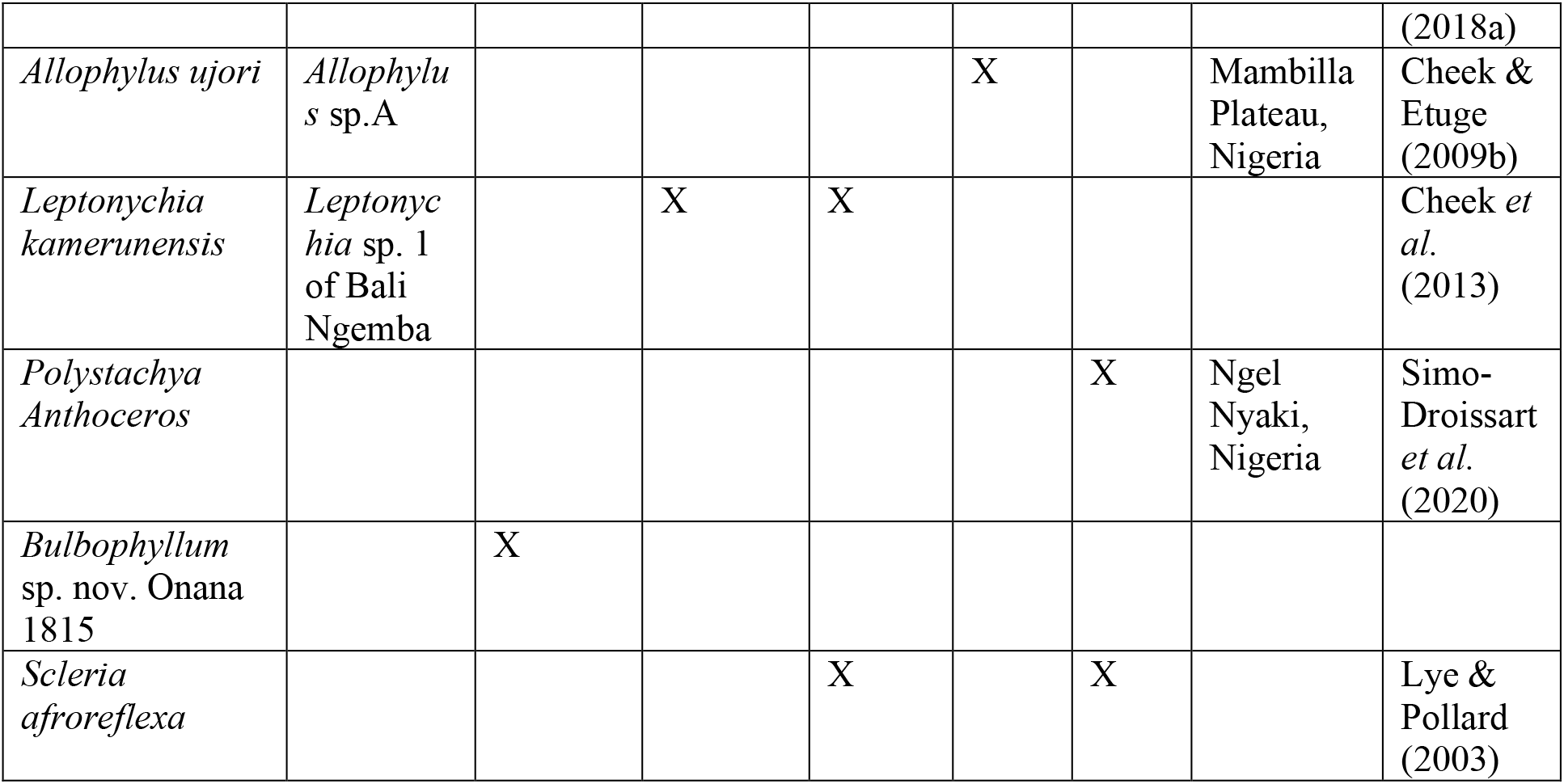
Bali Ngemba Forest Reserve, Cameroon: strictly endemic and near (also at one or two other locations). Revised and updated from Harvey *et al*. (2004: 49).

Examination of Table 2 shows that the number of strict endemics on Bali Ngemba is now nine, a drop of three from the 12 cited in Harvey *et al*. (2004). This change has partly resulted from further survey work in other locations which has extended the known ranges of the species concerned so that they are no longer strict endemics (Cheek *et al*. 2020b). It has also resulted from taxonomic revisions (Lachenaud 2019) and floristic studies (Couvreur *et al*. 2021) that have reduced to synonymy three of the previously supposed strict endemics.

The number of near-endemics for Bali Ngemba at 11, remains the same as in 2004. However, there has been some turnover with replacement on the list of formerly supposed near endemics since found to have wider ranges, with both former strict endemics now found to be near endemics and near endemic taxa that had not been listed as endemics in 2004.

By far the majority of near-endemics (5/11) of Bali Ngemba are shared with the Mt Kupe and the Bakossi Mts c. 120 km to the south along the Cameroon Highland Chain. Bali Ngemba represents the northern limit of these cloud forest species, excepting *Scleria afroreflexa* of montane grassland which extends north to Kilum-Ijim.

Yet Mt Kupe-Bakossi Mts have many endemics that do not extend to Bali Ngemba, e.g. *Impatiens frithii* Cheek (Cheek & Csiba 2002) and *Coffea montekupensis* Stoff. (Stoffelen *et al*. 1997), although the last does extend to Lebialem Highlands which is close to Bali Ngemba (Harvey *et al*. 2010).

While five of the provisionally named supposed endemics of Harvey *et al*. (2004) have now been formerly published (11 of which remain endemic or near-endemic) a further 12 (of which six are strictly endemic) remain unpublished. This paper continues the effort to address this deficit.

## Conclusions

About 2000 species of vascular plant have been described as new to science each year for the last fifteen years or more. Cameroon currently has the highest number of new species to science published each year (Cheek *et al*. 2020c), recent examples being published are (Achoundong *et al*. 2021; Aguirre-Alvarez *et al*. 2021; Cheek *et al*. 2021b; Cheek *et al*. 2021c; Cheek *et al*. 2021d; Couvreur *et al*. 2021; Gosline *et al*. 2021). Until species are known to science, they cannot be assessed for their conservation status and the possibility of protecting them is reduced (Cheek *et al*. 2020c). To maximise the survival prospects of range-restricted species such as *Vepris onanae*, there is an urgent need not only to document them in compliance with the nomenclatural code (Turland *et al*. 2018), but also to formally assess the species for their extinction risk rapidly, applying the criteria of a recognised system, of which the IUCN Red List of Threatened Species is the most universally accepted (Bachman *et al*. 2019). Thanks to the Global Tree Assessment (BGCI 2021) many of the world’s tree species have now been assessed. The State of the World’s Trees concluded that the highest proportion of threatened tree species is found in Tropical Africa, and that Cameroon has the highest number (414) of threatened tree species of all tropical African countries (BGCI 2021). However, the vast majority of plant species still lack assessments on the Red List (Nic Lughadha *et al*. 2020).

A demonstration that governments and leaders recognise the importance of species assessed as threatened by on the Red List, was seen recently in Cameroon when in part due to the high number of plant species on the Red List (Lovell 2020), a logging concession was revoked for the Ebo forest, a Tropical Important Plant Area or TIPA (Kew Science News 2020).

Thus, while the Conservation Status statement included in the species treatment in the present paper highlight the threatened status of *Vepris onanae* to the taxonomic botany community, completion and review of full assessments must follow so that it can be published on the IUCN Red List. Assessment on the Red List will facilitate inclusion in larger scale studies, such as future global extinction risk estimates and conservation prioritisation exercises, but, most importantly, it will enable not only scientists, but local communities, NGOs and national authorities to take action to safeguard them.

Documented extinctions of plant species are increasing (Humphreys *et al*. 2019) and recent estimates suggest that as many as two fifths of the world’s plant species are now threatened with extinction (Nic Lughadha *et al*. 2020). In Cameroon, *Oxygyne triandra* Schltr. and *Afrothismia pachyantha* Schltr. of South West Region, Cameroon are now known to be globally extinct (Cheek & Williams 1999, Cheek *et al*. 2018c, Cheek *et al*. 2019b). In some cases, Cameroon species appear to have become extinct even before they are known to science, such as *Vepris bali* Cheek which formerly grew and was endemic to Bali Ngemba together with (but at a different altitude to) *Vepris onanae* (Cheek *et al*. 2018a), and this is also true in neighbouring Gabon. e.g. *Pseudohydrosme bogneri* Cheek (Moxon-Holt & Cheek 2021).

Most of the 815 Cameroonian species in the Red Data Book for the plants of Cameroon are threatened with extinction due to habitat clearance or degradation, especially of forest for small-holder and plantation agriculture following logging (Onana & Cheek, 2011) while others are specifically targeted (e.g. Nana et al. 2021). Efforts are now being made to delimit the highest priority areas in Cameroon for plant conservation as Tropical Important Plant Areas (TIPAs) using the revised IPA criteria set out in Darbyshire *et al*. (2017). This is to help avoid the global extinction of additional endemic species, such as *Vepris onanae* published in this paper, which it is intended will be included in the proposed TIPAs of Mt Kupe and Bali Ngemba.

Bali Ngemba, while a Forest reserve, is not formally protected for nature conservation and is its natural forest habitat is being encroached for agriculture. Protection of Bali Ngemba, with the support of local communities is essential if the endemic plant species unique to it are not to become extinct globally, and if the 34 threatened species there are not to move closer to extinction.

## Acknowledgements

This paper was completed as part of the Cameroon TIPAs (Tropical Important Plant Areas) project at RBG, Kew, which is supported by Players of Peoples Postcode Lottery. Formal Red Listing of *Vepris onanae*, once this paper is formally published, will be supported by the John S. Cohen Foundation. We thank Lydia Burns and Penny Applebe of Kew’s Foundation for making this possible. The specimens cited in this paper were collected by or with the support of volunteers and sponsored scientists arranged by Earthwatch Europe, Oxford, and we were also assisted greatly by our colleagues such as the late Martin Etuge, but also Innocent Wultoff and Terence Atem. John DeMarco of the then Bamenda Highlands Forest Project persuaded us to make a botanical survey of Bali Ngemba and of Baba 2 forests in the Bamenda Highlands, and with Anne Gardner supported our visits. George Gosline and Stuart Cable of Royal Botanic Gardens, Kew gave invaluable support at Mt Kupe as did Chris and Liz Bowden of the former Mt Kupe Forest Project (Birdlife International). Drs Benoît Satabié, Gaston Achoundong and more recently Florence Ngo Ngwe, Eric Nana, Jean Lagarde, the current and former directors, of IRAD-National Herbarium of Cameroon, Yaoundé, and Betti their staff are thanked for expediting the collaboration between our two institutes. Janis Shillito typed the manuscript. Two anonymous reviewers are thanked for constructively reviewing an earlier version of this paper.

## References

Achoundong, G., van der Burgt, X., Cheek, M. (2021). Four new threatened species of *Rinorea* (Violaceae), treelets from the forests of Cameroon bioRxiv. https://doi.org/10.1101/2021.05.19.444792

Alvarez-Aguirre, M.G, Cheek, M., Sonké, B. (2021). *Kupeantha yabassi* (Coffeeae-Rubiaceae), a new Critically Endangered shrub species of the Ebo Forest area, Littoral Region, Cameroon. biorxiv https://doi.org/10.1101/2021.03.21.436301

Appelhans, M.S., Wen, J. (2020). Phylogenetic placement of *Ivodea* and biogeographic affinities of Malagasy Rutaceae. Plant Systematics and Evolution 306:1–14.

Atangana, A.F., Toze, F.A.A., Langat, M.K., Happi, N.H., Mbaze, L.L.M., Mulholland, D.A., et al. (2017). Acridone alkaloids from *Vepris verdoorniana* (Excell & Mendonça) Mziray (Rutaceae). Phytochemistry Letters 19:191–5.

Bachman, S.P., Field, R., Reader, T., Raimondo, D., Donaldson, J., Schatz, G.E. and Lughadha, E.N. (2019). Progress, challenges and opportunities for Red Listing. Biological Conservation 234: 45–55. https://doi.org/10.1016/j.biocon.2019.03.002

Bachman, S., Moat, J., Hill, A.W., de la Torre, J., Scott, B. (2011). Supporting Red List threat assessments with GeoCAT: geospatial conservation assessment tool, in: Smith V, Penev, eds. e-Infrastructures for data publishing in biodiversity science. ZooKeys 150:117–126. Available from: http://geocat.kew.org/ [accessed 17 Sept. 2021].

Barthlott, W., Lauer, W. & Placke, A. (1996). Global distribution of species diversity in vascular plants: towards a world map of phytodiversity. Erdkunde 50: 317–327 https://doi.org/10.1007/s004250050096

BGCI (2021). State of the World’s Trees. BGCI, Richmond, UK.

Breteler, F.J., Bakker, F.T. & Jongkind, C.C. (2015). A synopsis of *Soyauxia* (Peridiscaceae, formerly Medusandraceae) with a new species from Liberia. Plant Ecology and Evolution. 148: 409–419. https://doi.org/10.5091/plecevo.2015.1040

Burkill, H.M. (1997). The Useful Plants of West Tropical Africa. Vol. 4, families M-R. Royal Botanic Gardens, Kew.

Cable, S. & Cheek, M. (1998). The Plants of Mt Cameroon, a Conservation Checklist. Royal Botanic Gardens, Kew.

Chapman, J. & Chapman, H. (2001). The Forests of Taraba and Adamawa States, Nigeria an Ecological Account and Plant Species Checklist. University of Canterbury: Christchurch, New Zealand. pp. 221.

Cheek, M. (1992). A Botanical Inventory of the Mabeta-Moliwe Forest. Royal Botanic Gardens, Kew.

Cheek, M. (2003). A new species of *Ledermanniella (Podostemaceae)* from western Cameroon. Kew Bull. 58: 733–737. https://doi.org/10.2307/4111153

Cheek, M. (2004). Vepris lecomteana. The IUCN Red List of Threatened Species 2004: e.T46174A11039677. https://dx.doi.org/10.2305/IUCN.UK.2004.RLTS.T46174A11039677.en. (Downloaded on 22 May 2021).

Cheek, M. (2017). Vepris felicis. The IUCN Red List of Threatened Species 2017: e.T65064584A65064590. http://dx.doi.org/10.2305/IUCN.UK.2017-3.RLTS.T65064584A65064590.en. (accessed: 05/2021).

Cheek, M. & Cable, S. (1997). Plant Inventory for conservation management: the Kew-Earthwatch programme in Western Cameroon, 1993-96, pp. 29–38 in Doolan, S. (Ed.) African Rainforests and the Conservation of Biodiversity, Earthwatch Europe, Oxford.

Cheek, M. & Csiba, L. (2002). A new epiphytic species of *Impatiens* (Balsaminaceae) from western Cameroon. Kew Bull. 57: 669–674. https://doi.org/10.2307/4110997

Cheek, M. & Etuge, M. (2009a). A new submontane species of *Deinbollia* (Sapindaceae) from Western Cameroon and adjoining Nigeria. Kew Bull. 64: 503–508. https://doi.org/10.1007/s12225-009-9132-4

Cheek M. & Etuge, M. (2009b). *Allophylus conraui* (*Sapindaceae*) reassessed and *Allophylus ujori* described from western Cameroon. Kew Bull. 64(3): 495–502. https://doi.org/10.1007/s12225-009-9139-x

Cheek, M. & Onana, J-M. (2021). The endemic plant species of Mt Kupe, Cameroon with a new Critically Endangered cloud-forest tree species, *Vepris zapfackii* (Rutaceae). biorxiv. (pre-print) https://doi.org/10.1101/2021.06.01.446645

Cheek, M., Williams, S. (1999). A Review of African Saprophytic Flowering Plants. In: Timberlake, Kativu eds. African Plants. Biodiversity, Taxonomy & Uses. Proceedings of the 15th AETFAT Congress at Harare. Zimbabwe, 39–49.

Cheek, M., Achoundong, G., Onana, J-M., Pollard, B., Gosline, G., Moat, J., Harvey, Y.B. (2006). Conservation of the Plant Diversity of Western Cameroon. In: Ghazanfar SA, H.J. Beentje (eds). Proceedings of the 17th AETFAT Congress, Addis Ababa. Ethiopia, 779–791.

Cheek, M., Alvarez-Agiurre, M.G., Grall, A., Sonké, B., Howes, M-J.R., Larridon, L. (2018b). *Kupeantha* (Coffeeae, Rubiaceae), a new genus from Cameroon and Equatorial Guinea. PLoS ONE 13: 20199324. https://doi.org/10.1371/journal.pone.0199324

Cheek, M., Arcate, J., Choung, H.S., Herian, K., Corcoran, M., Horwath, A. (2013). Three new or resurrected species of *Leptonychia (Sterculiaceae-Byttneriaceae-Malvaceae)* from West-Central Africa. Kew Bull. 68. http://dx.doi.org/10.1007/s12225-013-9469-6)

Cheek, M., Cable, S., Hepper, F.N., Ndam, N., Watts, J. (1996). Mapping plant biodiversity on Mt. Cameroon. In: Maesen, Burgt, Rooy eds. The Biodiversity of African Plants (Proceedings XIV AETFAT Congress. Cameroon: Kluwer, 110–120. https://doi.org/10.1007/978-94-009-0285-5_16

Cheek, M., Causon, I., Tchiengue, B. & House, E. (2020b). Notes on the endemic cloud forest plants of the Cameroon Highlands and the new, Endangered, *Tricalysia elmar* (Coffeeae-Rubiaceae). Plant Ecology and Evolution 153: 167–176 https://doi.org/10.5091/plecevo.2020.1661

Cheek, M., Corcoran, M. & Horwath, A. (2009b). Four new submontane species of *Psychotria* (*Rubiaceae*) with bacterial nodules from western Cameroon. Kew Bull. 63: 405–418. https://doi.org/10.1007/s12225-008-9056-4

Cheek, M., Etuge, M. & Williams, S. (2019b). *Afrothismia kupensis* sp. nov. (Thismiaceae), Critically Endangered, with observations on its pollination and notes on the endemics of Mt Kupe, Cameroon. Blumea 64: 158–164.

Cheek, M., Gosline, G. & Onana, J.M. (2018a). *Vepris bali* (Rutaceae), a new critically endangered (possibly extinct) cloud forest tree species from Bali Ngemba, Cameroon. Willdenowia 48: 285–292. https://doi.org/10.3372/wi.48.48207

Cheek, M., Harvey, Y.B., Onana, J-M. (2010). The Plants of Dom. Bamenda Highlands, Cameroon: A Conservation Checklist. Royal Botanic Gardens, Kew.

Cheek M, Harvey Y, Onana J-M. (2011). The Plants of Mefou Proposed National Park. Yaoundé, Cameroon: A Conservation Checklist. Royal Botanic Gardens, Kew.

Cheek, M., Mackinder, B., Gosline, G., Onana, J.M., Achoundong, G. (2001). The phytogeography and flora of western Cameroon and the Cross River-Sanaga River interval. Systematics and Geography of Plants 71: 1097–1100. https://doi.org/10.2307/3668742

Cheek, M., Molmou, D., Magassouba, S., Ghogue, J.-P. (2021b). Taxonomic Monograph of *Saxicolella* (Podostemaceae), African waterfall plants highly threatened by Hydro-Electric projects, with five new species. bioRxiv: https://doi.org/10.1101/2021.06.19.449102

Cheek, M., Ndam, N., & Budden, A. (2021c). Notes on the threatened lowland forests of Mt Cameroon and their endemics including *Drypetes burnleyae* sp. nov., with a key to species of Drypetes sect. Stipulares (Putranjivaceae). Kew Bulletin, 1–12. https://doi.org/10.1007/s12225-021-09947-2

Cheek, M., Nic Lughadha, E., Kirk, P., Lindon, H., Carretero, J., Looney, B., Douglas, B., Haelewaters, D., Gaya, E., Llewellyn, T., Ainsworth, M., Gafforov, Y., Hyde, K., Crous, P., Hughes, M., Walker, B.E., Forzza, R.C., Wong, K.M., Niskanen, T. (2020c). New scientific discoveries: plants and fungi. Plants, People Planet 2: 371–388. https://doi.org/10.1002/ppp3.10148

Cheek, M., Oben, B. & Heller, T. (2009a). The identity of the West-Central African *Oricia lecomteana* Pierre, with a new combination in *Vepris* (Rutaceae). Kew Bull. 64: 509–512 https://doi.org/10.1007/s12225-009-9135-1

Cheek, M., Onana, J.M., Chapman, H.M. (2021a). The montane trees of the Cameroon Highlands, West-Central Africa, with *Deinbollia onanae* sp. nov. (Sapindaceae), a new primate-dispersed, Endangered species. PeerJ 9:e11036 https://doi.org/10.7717/peerj.11036

Cheek, M., Onana, J-M., Pollard, B.J. (2000). The Plants of Mount Oku and the Ijim Ridge, Cameroon, a Conservation Checklist. Royal Botanic Gardens, Kew.

Cheek, M., Onana, J-M., Yasuda, S., Lawrence, P., Ameka, G., Buinovskaja G. (2019a). Addressing the *Vepris verdoorniana* complex (Rutaceae) in West Africa, with two new species. Kew Bull. 74: 53. https://doi.org/10.1007/S12225-019-9837-Y

Cheek, M., Pollard, B.J., Darbyshire, I, Onana, J.M. & Wild, C. (2004). The Plants of Kupe, Mwanenguba and the Bakossi Mts, Cameroon. A Conservation Checklist. Royal Botanic Gardens, Kew.

Cheek, M., Satabie, B. & J-M. Onana (1997). Interim report on botanical survey and inventory for Kilum and Ijim Mountain Forest Projects by the National Herbarium of Cameroon and R.B.G., Kew Oct./Nov. 1996.

Cheek, M., Tchiengue, B., Baldwin, I. (2020a). Notes on the plants of Bakossi, Cameroon, and the new *Cola etugei* and *Cola kodminensis* (Sterculiaceae-Malvaceae). Plant Ecology and Evolution 153: 108–119 https://doi.org/10.5091/plecevo.2020.1662

Cheek, M., Tchiengué, B., van der Burgt, X. (2021d). Taxonomic revision of the threatened African genus *Pseudohydrosme* Engl. (Araceae), with *P. ebo*, a new, critically endangered species from Ebo, Cameroon. PeerJ 9:e10689 https://doi.org/10.7717/peerj.10689.

Cheek, M., Tsukaya, H., Rudall, P.J., Suetsugu, K. (2018c). Taxonomic monograph of Oxygyne (Thismiaceae), rare achlorophyllous mycoheterotrophs with strongly disjunct distribution. PeerJ 6: e4828. https://doi.org/10.7717/peerj.4828

Cheplogoi, P.K., Mulholland, D.A., Coombes, P.H., Randrianarivelojosia, M. (2008). An azole, an amide and a limonoid from *Vepris uguenensis* (Rutaceae). Phytochemistry 69: 1384–1388. https://doi.org/10.1016/j.phytochem.2007.12.013

Collar, N.J. & Stuart, S.N. (1988). Mount Oku (Cameroon) pp. 29–30 in Key Forests for Threatened Birds in Africa. ICBP Monograph No. 3. IUCN.

Courade, G. (1974). Commentaire des Cartes. Atlas Regional. Ouest 1. ORSTOM. Yaoundé, Cameroun.

Couvreur, T.L.P., Dagallier, L.-P.M.J., Crozier, F., Ghogue, J.-P., Hoekstra, P.H., Kamdem, N.G., Johnson, D.M., Murray, N., Sonké, B. (2021). Flora of Cameroon – Annonaceae. Phytokeys

Dagallier, L.P., Janssens, S.B., Dauby, G., Blach-Overgaard, A., Mackinder, B.A., Droissart, V., Svenning, J.C., Sosef, M.S., Stévart, T., Harris, D.J. & Sonké, B. (2020). Cradles and museums of generic plant diversity across tropical Africa. New Phytologist 225: 2196–2213.

Darbyshire, I., Anderson, S., Asatryan, A., Byfield, A., Cheek, M., Clubbe, C., Ghrabi, Z., Harris, T., Heatubun, C. D., Kalema, J., Magassouba, S., McCarthy, B., Milliken, W., Montmollin, B. de, Nic Lughadha, E., Onana, J.M., Saidou, D., Sarbu, A., Shrestha, K. & Radford, E. A. (2017). Important Plant Areas: revised selection criteria for a global approach to plant conservation. Biodivers. Conserv. 26: 1767–1800. https://doi.org/10.1007/s10531-017-1336-6.

Gereau, R.E. (2001). New names in African Celastraceae and Rutaceae. Novon 11:43–44.

Gosline, G. (2009). *Diospyros korupensis* sp. nov. and *Diospyros onanae* sp. nov.(Ebenaceae) from Cameroon. Nordic Journal of Botany. 27(5):353–358.

Gosline, G., Cheek, M., Onana, J-M., Ngansop, E., van der Burgt, X. & Dagallier, L.-P. (2021). *Uvariopsis ebo* (Annonaceae) a new, Critically Endangered tree species from the Ebo Forest, Cameroon and a key to the Cameroonian species of *Uvariopsis*. bioRxiv. https://doi.org/10.1101/2021.03.26.437154

Harris, D.J. (2000). Validation of the name *Vepris glaberrima* (Rutaceae). Kew Bulletin 55(2):458–458.

Harvey, Y.B., Pollard, B.J., Darbyshire, I., Onana, J.-M., Cheek, M. (2004). The Plants of Bali Ngemba Forest Reserve. Cameroon: A Conservation Checklist. Royal Botanic Gardens, Kew.

Harvey, Y.B., Tchiengue, B., Cheek, M. (2010). The Plants of the Lebialem Highlands, a Conservation Checklist. Royal Botanic Gardens, Kew.

Hawkins, P. & Brunt, M. (1965). The Soils and Ecolgy of West Cameroon. 2 vols. FAO, Rome. 516 pp.

Humphreys, A.M., Govaerts, R., Ficinski, S.Z., Lughadha, E.N. and Vorontsova, M.S. (2019). Global dataset shows geography and life form predict modern plant extinction and rediscovery. Nature Ecology & Evolution 3.7: 1043–1047. https://doi.org/10.1038/s41559-019-0906-2

Imbenzi, P.S., Osoro, E.K., Aboud, N.S., Ombito, J.O., Cheplogoi, P.K. (2014). A review on chemistry of some species of genus *Vepris* (*Rutaceae* family). Journal of Scientific and Innovative Research 3: 357–362

IPNI (continuously updated). The International Plant Names Index. http://ipni.org/ (accessed: 05/2020).

IUCN. (2012). IUCN Red List Categories and Criteria: Version 3.1. Second edition. - Gland, Switzerland and Cambridge, UK: IUCN. Available from: http://www.iucnredlist.org/ (accessed: May 2021).

Kew Science News (2020). Ebo Forest Logging Plans Suspended. https://www.kew.org/read-and-watch/ebo-forest-logging-suspended (accessed 5 May 2021).

Lachenaud, O. (2019). Révision du Genre *Psychotria* (Rubiaceae) en Afrique Occidentale et Centrale. Opera Botanica Belgica 17: 1–900.

Lachenaud, O. & Onana J. M. (2021). The West and Central African species of *Vepris* Comm. ex A.Juss. (Rutaceae) with simple or unifoliolate leaves, including two new combinations. Adansonia 43: 107–116. https://doi.org/10.5252/adansonia2021v43a10. http://adansonia.com/43/10

Langat, M.K. (2011). Flindersiamine, a Furoquinoline Alkaloid from *Vepris uguenensis* (Rutaceae) as a Synergist to Pyrethrins for the Control of the Housefly, *Musca domestica* L. (Diptera: Muscidae). Journal of the Kenya Chemical Society. 6: 9–15.

Langat, M., Kami, T., Cheek, M. (2021a). Chemistry, Taxonomy and Ecology of the potentially chimpanzee-dispersed *Vepris teva* sp.nov. (Rutaceae) of coastal thicket in the Congo Republic. biorxiv. https://doi.org/10.1101/2021.08.22.457282

Langat, M.K., Mayowa, Y., Sadgrove, N., Danyaal, M., Prescott, T.A., Kami, T., Schwikkard, S., Barker, J. & Cheek, M. (2021b). Multi-layered antimicrobial synergism of (E)-caryophyllene with minor compounds, tecleanatalensine B and normelicopine, from the leaves of *Vepris gossweileri* (I. Verd.) Mziray. Natural Product Research, pp.1–11. https://doi.org/10.1080/14786419.2021.1899176

Letouzey, R. (1963). Rutaceae. Flore du Cameroun 1. Muséum Nationale D’Histoire Naturelle, Paris.

Lovell, R. (2020). Timber! The threat to Cameroon’s Ebo Forest. https://www.kew.org/read-and-watch/ebo-forest-cameroon (accessed 24 Sept. 2021).

Lovell, R. & Cheek, M. 2021. Vepris adamaouae. The IUCN Red List of Threatened Species 2021: e.T190265400A190322551. https://dx.doi.org/10.2305/IUCN.UK.2021-2.RLTS.T190265400A190322551.en. Downloaded on 24 September 2021.

Lye, K. A., & Pollard, B. J. (2003). Studies in African Cyperaceae 29. *Scleria afroreflexa*, a new species from western Cameroon. Nord. J. Bot. 23(4): 431–435.

Maisels, F.M., Cheek, M., Wild, C. (2000). Rare plants on Mt Oku summit, Cameroon. Oryx 34: 136–140. https://doi.org/10.1017/s0030605300031057.

Morton, C.M. (2017). Phylogenetic relationships of *Vepris (Rutaceae)* inferred from chloroplast, nuclear, and morphological data. PLoS ONE 12:e0172708. https://doi.org/10.1371/journal.pone.0172708

Moxon-Holt, L. & Cheek, M. (2020). *Pseudohydrosme bogneri* sp. nov. (Araceae), a spectacular Critically Endangered (Possibly Extinct) species from Gabon, long confused with *Anchomanes nigritianus*. BioRxiv (pre-print) https://doi.org/10.1101/2021.03.25.437040

Mziray, W. (1992). Taxonomic studies in *Toddalieae* Hook.f. (*Rutaceae*) in Africa. Symb. Bot. Upsal. 30: 1–95.

Nana, E., Takor, C., Nkengbeza, S., Tchiengue, B., Ngansop, E., & Tchopwe, E. (2021). Illegal logging threatens to wipe out the Critically Endangered African zebrawood Microberlinia bisulcata from Cameroon’s Ebo forest. Oryx, 55(5), 652–653. doi:10.1017/S0030605321000910

Nic Lughadha, E., Bachman, S.P., Leão, T.C., Forest, F., Halley, J.M., Moat, J., Acedo, C., Bacon, K.L., Brewer, R.F., Gâteblé, G., Gonçalves, S.C., Govaerts, R., Hollingsworth, P.M., Krisai-Greilhuber, I., de Lirio, E.J., Moore, P.G.P., Negrão, R., Onana, J.M., Rajaovelona, L.R., Razanajatovo, H., Reich, P.B., Richards, S.L., Rivers, M.C., Amanda Cooper, A., Iganci, J., Lewis, G.L., Smidt, E.C., Antonelli, A., Mueller, G.M. & Walker, B.E. (2020). Extinction risk and threats to plants and fungi. Plants, People, Planet 2: 389 –408. https://doi.org/10.1098/rstb.2017.0402

Ombito, J.O., Chi, G.F. & Wansi, J.D. (2020). Ethnomedicinal uses, phytochemistry, and pharmacology of the genus Vepris (Rutaceae): A review. Journal of Ethnopharmacology, p.113622. https://doi.org/10.1016/j.jep.2020.113622

Onana, J.M. (2011). The Vascular Plants of Cameroon. A taxonomic checklist with IUCN *assessments*. Flore du Cameroun 39”occasional volume “. IRAD-National Herbarium of Cameroun.

Onana, J.M.(2013). Synopsis des espèces végétales vasculaires endémiques et rares du Cameroun. Check-liste pour la conservation de la biodiversité. Flore du Cameroun 40. Ministère de la Recherche Scientifique et de l’Innovation (MINRESI), Yaoundé, Cameroun.

Onana, J.M. (2017). Burseraceae. Flore du Cameroun 43. Ministère de la Recherche Scientifique et de l’Innovation (MINRESI), Yaoundé, Cameroun

Onana, J.M. & Cheek, M. (2011). Red Data Book of the Flowering Plants of Cameroon, IUCN Global Assessments. Royal Botanic Gardens, Kew.

Onana, J.M. & Chevillotte, H. (2015). Taxonomie des *Rutaceae – Toddalieae* du Cameroun revisitée: découverte de quatre espèces nouvelles, validation d’une combinaison nouvelle et véritable identité de deux autres espèces de *Vepris* Comm. ex A. Juss. Adansonia, sér. 3. 37: 103–129. https://doi.org/c10.5252/a2015n1a7

Onana, J. M., Cheek, M., & Chevillotte, H. (2019). Additions au genre *Vepris* Comm. ex A. Juss. (Rutaceae-Toddalieae) au Cameroun. Adansonia 41: 41–52. https://doi.org/10.5252/adansonia2019v41a5

Plants of the World Online (continuously updated). Facilitated by the Royal Botanic Gardens, Kew. http://www.plantsoftheworldonline.org/?f=accepted_names&q=Vepris (downloaded 1 May 2021)

Shorthouse, D.P. (2010). SimpleMappr, an online tool to produce publication-quality point maps. [Retrieved from http://www.simplemappr.net accessed 28 May 2021]

Simo-Droissart, M., Stévart, T., Pollard, B.J. & Droissart, V. (2020). Polystachya anthoceros. The IUCN Red List of Threatened Species 2020:e.T87751188A87757909. https://dx.doi.org/10.2305/IUCN.UK.2020-3.RLTS.T87751188A87757909.en. Downloaded on 22 September 2021.

Soltis, D.E., Clayton, J.W., Davis, C.C., Wurdack, K.J., Gitzendanner, M.A., Cheek, M., Savolainen, V., Amorim, A.M. & Soltis, P.S. (2007). Monophyly and relationships of the enigmatic family *Peridiscaceae*. Taxon 56: 65–73.

Sosef, M.S.M., Wieringa, J.J., Jongkind, C.C.H., Achoundong, G., Azizet Issembé, Y., Bedigian, D., Van Den Berg, R.G., Breteler, F.J., Cheek, M., Degreef, J. (2006). Checklist of Gabonese Vascular Plants. Scripta Botanica Belgica 35. National Botanic Garden of Belgium.

Stoffelen, P., Cheek, M., Bridson, D., Robbrecht. E. (1997). A new species of *Coffea* (Rubiaceae) and notes on Mt Kupe (Cameroon). Kew Bull. 52: 989 –994. https://doi.org/10.2307/3668527

Tame, S. & Asonganyi, J. (1995). Ijim Mountain Forest project Plant List. Appendix 2. 16 pp.

Thomas, D. W. (1986). Provisional Species List for Mount Oku Flora. Pp. 59-62 in McLeod, H. L. The Conservation of Oku Mountain Forest, Cameroon. Study report No. 15. 90 pp. ICBP.

Thiers, B. (continuously updated). Index Herbariorum: A global directory of public herbaria and associated staff. New York Botanical Garden’s Virtual Herbarium. http://sweetgum.nybg.org/ih/ (accessed: Jan. 2021).

Turland, N.J., Wiersema, J.H., Barrie, F. R., Greuter, W., Hawksworth, D.L., Herendeen, P.S., Knapp, S., Kusber, W.-H., Li, D.-Z., Marhold, K., May, T.W., McNeill, J., Monro, A.M., Prado, J., Price, M.J. & Smith, G.F. (ed.) (2018). International Code of Nomenclature for algae, fungi, and plants (Shenzhen Code) adopted by the Nineteenth International Botanical Congress Shenzhen, China, July 2017. – Glashütten: Koeltz Botanical Books. [= Regnum Veg. 159].

Verdoorn, I. C. (1926). Revision of African Toddalieae. Bulletin of Miscellaneous Information, Royal Botanic Gardens, Kew 9: 389 –416. https://www.jstor.org/stable/4118639

Wansi, J.D., Mesaik, M., Chiozem, D.D., Devkota, K.P., Gaboriaud-Kolar, N., Lallemand, M-C., Wandji, J., Choudhary, M.I., & Sewald, N. (2008). Oxidative Burst Inhibitory and Cytotoxic Indoloquinazoline and Furoquinoline Alkaloids from *Oricia suaveolens*. Journal of Natural Products 71: 1942 –1945. https://doi.org/10.1021/np800276f

Whittaker, A. & Cheek, M. 2021. Vepris montisbambutensis. The IUCN Red List of Threatened Species 2021:e.T190266398A190322631. https://dx.doi.org/10.2305/IUCN.UK.2021-2.RLTS.T190266398A190322631.en. Downloaded on 24 September 2021

World Conservation Monitoring Centre. (1998). Vepris trifoliolata. The IUCN Red List of Threatened Species 1998: e.T46175A11033334. https://dx.doi.org/10.2305/IUCN.UK.1998.RLTS.T46175A11033334.en

